# Robust and stable transcriptional repression in *Giardia* using CRISPRi

**DOI:** 10.1101/358598

**Authors:** SG McInally, KD Hagen, C Nosala, J Williams, K Nguyen, J Booker, K Jones, C. Dawson Scott

## Abstract

*Giardia lamblia* is a binucleate protistan parasite causing significant diarrheal disease worldwide. An inability to target Cas9 to both nuclei, combined with the lack of non-homologous end joining and markers for positive selection, has stalled the adaptation of CRISPR/Cas9-mediated genetic tools for this widespread parasite. CRISPR interference (CRISPRi) is a modification of the CRISPR/Cas9 system that directs catalytically inactive Cas9 (dCas9) to target loci for stable transcriptional repression. Using a *Giardia* nuclear localization signal to target dCas9 to both nuclei, we developed efficient and stable CRISPRi-mediated transcriptional repression of exogenous and endogenous genes in *Giardia*. Specifically, CRISPRi knockdown of kinesin-2a and kinesin-13 causes severe flagellar length defects that mirror defects with morpholino knockdown. Knockdown of the ventral disc MBP protein also causes severe structural defects that are highly prevalent and persist in the population more than five days longer than transient morpholino-based knockdown. By expressing two gRNAs in tandem to simultaneously knock down kinesin-13 and MBP, we created a stable dual knockdown strain with both flagellar length and disc defects. The efficiency and simplicity of CRISPRi in polyploid *Giardia* allows for rapid evaluation of knockdown phenotypes and highlights the utility of CRISPRi for emerging model systems.

## Introduction

*Giardia lamblia* is a common parasitic protist that infects over three hundred million people worldwide each year, causing acute diarrheal disease in areas with inadequate sanitation and water treatment (Einarsson et al., 2016). *Giardia* has a two-stage life cycle in which mammalian hosts ingest quadrinucleate cysts from contaminated water sources. As they pass into the gastrointestinal tract, cysts develop into binucleate, flagellated trophozoites. During infection, trophozoites attach extracellularly to the gut epithelium, proliferate, and later differentiate back into infectious cysts that are passed into the environment (Heyworth, 2014). Giardiasis is considered a neglected disease (Savioli et al., 2006), and the need for new effective and affordable treatments for giardiasis is underscored by estimated failure rates of up to 20% for standard treatments (Upcroft and Upcroft, 2001), emerging evidence of drug resistance (Barat and Bloland, 1997; Land and Johnson, 1999; Upcroft et al., 1996), and the prevalence of chronic or recurrent infections (Bartelt and Sartor, 2015).

The nuclei in trophozoites and cysts are diploid (Bernander et al., 2001), transcriptionally active (Kabnick and Peattie, 1990), and genetically equivalent (Adam et al., 2013; Franzen et al., 2009; Hanevik et al., 2015; Morrison et al., 2007). Because *Giardia* trophozoites are effectively tetraploid and lack a defined sexual cycle (Poxleitner et al., 2008), the development of molecular genetic tools in *Giardia* has generally lagged behind other parasitic protists. Although new CRISPR/Cas9 gene-editing strategies have revolutionized molecular genetics in other protists (Grzybek et al., 2018), the adaptation of CRISPR-based tools for *Giardia* has been hindered by *Giardia*’s lack of a non-homologous end-joining (NHEJ) pathway (Morrison et al., 2007) and by the inability to target native *Streptomyces pyogenes* Cas9 (SpCas9) to the two nuclei. The widely used SV40 nuclear localizing signal (NLS) failed to localize full-length Cas9 (Ebneter et al., 2016), although it has been used previously in *Giardia* to localize exogenously expressed GFP or TetR protein to the nuclei (Elmendorf et al., 2000). For these reasons, the first *Giardia* quadruple knockout strain of the cyst wall component CWP1 was constructed using Cre/loxP sequential gene disruption, rather than a CRISPR/Cas9-mediated strategy (Ebneter et al., 2016).

Gene knockouts are only one of many strategies to interrogate gene function (Qi et al., 2013), and null alleles in essential processes like cell cycle regulation (Horlock-Roberts et al., 2017) and cytokinesis (Hardin et al., 2017) can result in severely reduced fitness or lethality. Translational repression using morpholino oligonucleotides has been used extensively in *Giardia* (House et al., 2011; Paredez et al., 2011; Woessner and Dawson, 2012), but morpholinos are transient (lasting less than 48 hours), lack complete penetrance, and are costly, making morpholino knockdowns less useful for genome-wide functional screens (Krtkova and Paredez, 2017). Both the transience and incomplete penetrance of morpholino knockdowns are problematic for evaluation of infection dynamics of *Giardia* mutants (Barash, 2017). Transcriptional repression is an important alternative strategy for the characterization of gene function; however, despite the presence of conserved components of the RNAi machinery, RNAi does not efficiently silence genes in *Giardia* (Krtkova and Paredez, 2017).

CRISPR/Cas9 is a modular and flexible DNA-binding platform that has been adapted for applications beyond genome editing, including transcriptional repression (Larson et al., 2013). The CRISPR interference system (CRISPRi) is a modification of the CRISPR/Cas9 system, in which a catalytically inactive or “dead” Cas9 (dCas9) induces stable, inducible, or reversible gene knockdown in eukaryotes (Larson et al., 2013; Piatek et al., 2015), as well as diverse bacteria (Kaczmarzyk et al., 2018; Larson et al., 2013; Liu et al., 2017; Tao et al., 2017; Zhang et al., 2016; Zuberi et al., 2017). Using a complementary guide RNA (gRNA), the inactive dCas9 is directed to precise genomic targets, where it binds and inhibits transcription initiation and/or elongation rather than inducing double-stranded breaks in DNA (Larson et al., 2013). CRISPRi has significant advantages over RNAi or morpholinos as it directly and stably inhibits transcription (Larson et al., 2013). In many systems, CRISPRi is empirically as effective as RNAi in transcriptional silencing, with significantly fewer off-target effects (Larson et al., 2013).

Here we demonstrate precise and stable CRISPRi-mediated transcriptional repression of both exogenous and endogenous genes in *Giardia*. This first successful application of CRISPRi for transcriptional knockdown in a parasitic protist—or a polyploid eukaryote—highlights the utility of CRISPRi for emerging model systems. Using a *Giardia* nuclear localization signal (NLS) to target dCas9 to both nuclei, we show efficient and persistent CRISPRi-mediated repression of one exogenous reporter gene and three endogenous *Giardia* cytoskeletal genes. CRISPRi knockdowns have severe cytoskeletal phenotypes that are highly prevalent and persist at least one week in cultured trophozoites. We also tandemly express gRNAs to simultaneously knock down one flagellar and one ventral disc gene, resulting in a stable strain with two knockdown phenotypes. The efficiency and simplicity of CRISPRi in *Giardia* permits the rapid assessment of the phenotypic consequences of protein knockdown (Jost et al., 2017; Larson et al., 2013) and will likely enable the first forward and reverse genetic screens.

## Methods

### Construction of the expression vector Cas9-SV40NLS-GFP for Cas9 localization

The expression vector Cas9-SV40NLS-GFP was constructed by placing mammalian codon-optimized *Streptococcus pyogenes* Cas9 (mCas9) with a C-terminal SV40 nuclear localizing signal (NLS) from JDS246 (Addgene #43861, JK Joung, unpublished) under the control of the *Giardia* malate dehydrogenase (MDH; GiardiaDB GL50803_3331) promoter. A 3xHA epitope tag widely used in *Giardia* (Gourguechon and Cande, 2011) was added to the C-terminal end and the resulting P_MDH_-mCas9-SV40NLS-3HA fragment was cloned into our C-terminal GFP vector, pcGFP1Fpac (Hagen et al., 2011), fusing Cas9 to GFP.

### Identification of putative Giardia NLSs and construction of vectors to test NLS efficacy

Putative NLS sequences were identified among the protein sequences of 65 *Giardia* strains expressing C-terminal GFP-tagged proteins that localize to the nuclei (Aurrecoechea et al., 2009) using the NLS prediction software NLStradamus (Nguyen Ba et al., 2009), with the two state HMM static model and posterior prediction with a cutoff of 0.6. To delete C-terminal NLSs, we amplified protein coding regions plus 200 bp of upstream sequence to include native promoters using reverse primers that excluded the NLSs and any downstream sequence. The truncated fragments were cloned into pcGFP1Fpac (Hagen et al., 2011). NLS sequences were added to Cas9 either by replacing the GFP tag in Cas9-SV40NLS-GFP with the NLS sequence to be tested (for C-terminal NLSs), or by adding a new start codon and NLS in front of the start codon of Cas9 (for N-terminal NLSs). NLS sequences were generated by PCR amplification from genomic DNA or by annealing short complementary oligonucleotides.

### Construction of dCas9 CRISPR interference vector dCas9g1pac

dCas9g1pac was created by amplifying the GL50803_2340 NLS from *Giardia* genomic DNA with 2340NLSAgeF (5’-actgct<u>accggt</u>cctcccagagaagaagcggtccaag-3’) and 2340NLSNotIR (5’- actgct<u>gcggccgc</u>tttagctcttaattttactaactctacgatcc-3’) and cloning it into Cas9-SV40NLS-GFP to replace GFP. A gRNA expression cassette with inverted BbsI sites for cloning specific gRNA targeting sequences was added, followed by the gRNA scaffold sequence from pX330 (Cong et al., 2013). To drive gRNA expression in *Giardia* and ensure proper termination, we included approximately 200 bp of DNA located upstream and downstream of the *Giardia* U6 spliceosomal snRNA (Hudson et al., 2012). The upstream sequence includes 12 bp of U6 coding sequence, and the downstream sequence includes the 12 bp motif thought to mediate 3’ end processing (Hudson et al., 2012). The gRNA cassette was synthesized (Biomatik) and inserted into a unique Acc65I site in the vector. The dCas9 mutations D10A and H840A were added by replacing a 3.6 kb NcoI-NheI fragment within Cas9 with a similar fragment from dCas9 vector MSP712 (Addgene #65768) (Kleinstiver et al., 2015). As constructed, the dCas9g1pac vector contains an 18 bp non-specific sequence between the U6 promoter and gRNA scaffold sequence that also includes the inverted BbsI sites for cloning annealed guide oligos. All BLAST hits for this sequence to the *Giardia* ATCC 50803 genome lack the protospacer adjacent motif (PAM) that is essential for CRISPRi silencing (Larson et al., 2013); all also have mismatches in the critical ‘seed region’ required for dCas9 binding (Qi et al., 2013) or have six or more mismatches throughout the guide sequence. Trophozoites carrying the ‘non-specific gRNA’ vector have no discernable phenotype.

### Guide RNA design and cloning

Specific guide RNAs (20 nt) were designed with the CRISPR ‘Design and Analyze Guides’ tool from Benchling (https://benchling.com/crispr) using a NGG PAM sequence and the *Giardia lamblia* ATCC 50803 genome (GenBank Assembly GCA_000002435.1). gRNAs were designed to target the non-template strand unless otherwise specified (Supplemental Material). gRNA oligonucleotides with 4-base overhangs complementary to the vector sequence overhangs were annealed and cloned into BbsI-digested dCas9g1pac. To express two gRNAs, each was individually cloned and then a universal primer set (gRNA_multiplex_F: 5’-gtcttataaggtaccgagcttgattgcaatagcaaacag-3’ and gRNA_multiplex_R: 5’-ctatagggcgaattcgagctgagctcggtaccttataagacaacatc-3’) was used to amplify the entire gRNA cassette from one plasmid and insert it into the SacI site of the second via Gibson assembly (Gibson et al., 2009). This method was also used to generate the non-specific gRNA plasmid dCas9g2pac, which has two gRNA cassettes, each with the non-specific gRNA sequence used in dCas9g1pac. While we have only used two gRNAs here, this method regenerates the SacI restriction site, allowing the insertion of additional gRNAs using the same primers (Supplemental Material).

### Construction of the NanoLucNeo luciferase expression vector

The NanoLucNeo vector was constructed via PCR amplification of the MDH promoter (GiardiaDB GL50803_3331), and NanoLuc (pFN31K, Promega). The resulting amplicons were cloned via Gibson assembly into BamHI and EcoRI digested pKS_mNeonGreen-N11_NEO (Hardin et al., 2017).

### Strains and culture conditions

All *G. lamblia* (ATCC 50803) strains were cultured in modified TYI-S-33 medium supplemented with bovine bile and 5% adult and 5% fetal bovine serum [56] in sterile 16 ml screw-capped disposable tubes (BD Falcon), and incubated upright at 37°C without shaking. Vectors were introduced into WBC6 or NanoLucNeo trophozoites by electroporation (~20 µg DNA) as previously described (Hagen et al., 2011). Strains were maintained with antibiotic selection (50 µg/ml puromycin and/or 600 µg/ml G418) (Hagen et al., 2011). All CRISPRi strains were thawed from frozen stocks and cultured for 24 to 48 hours prior to phenotypic analysis, unless otherwise specified.

### In vitro bioluminescence assays of NanoLuc knockdown strains

*Giardia* trophozoites were grown to confluency at 37°C for 48 hours, iced for 15 minutes to detach trophozoites and centrifuged at 900 x g for five minutes at 4°C. Trophozoites were resuspended in 1 ml of cold media and serial dilutions (10^-^1, 10^-^2) were made for enumeration using a hemocytometer. To measure luminescence, three 50 µl aliquots of the 10^-^2 dilution (approximately 10^3^ cells) were loaded into a white opaque 96-well assay plate (Corning Costar) and incubated for 30 minutes at 37°C. Following incubation, 50 µl of Nano-Glo^®^ Luciferase assay reagent (Promega), prepared at a 1:50 ratio of substrate to buffer, was added to each well. Luminescence was analyzed on a VictorX3 plate reader warmed to 37°C, using 0.1-second exposures, repeated every 30 seconds until maximal signal was detected. Experiments were performed using two independent samples with three technical replicates that were averaged over three different luminescence acquisitions and displayed with 95% confidence intervals. To determine the linear dynamic range of detection for the NanoLucNeo strain, we used tenfold serial dilutions and plotted the luminescence readings versus the number of cells loaded into each well (Supplemental Material).

### Quantitation of transcriptional knockdown using quantitative PCR (qPCR)

RNA was extracted using a commercial kit (Zymo Research). RNA quality was assessed by spectrophotometric analysis and electrophoresis prior to double-stranded cDNA synthesis using the QuantiTect Reverse Transcription Kit (Qiagen). Quantitative PCR of kinesin-2a (GiardiaDB GL50803_17333) was performed using 17333_qPCR1F 5’ GCCTCAACCAACTACGACGA 3’ and 17333_qPCR1R 5’ TCAGCACATCCATCGGCTTT 3’. Quantitative PCR of kinesin-13 (GiardiaDB GL50803_16945) was performed with 16945_qPCR3F 5’ CCAATACGCTGCAAAGCCTC 3’ and 16945_qPCR3R 5’ AGCCAGTTTGTCCATACGCA 3’. Analyses were performed using SensiFast No-ROX SYBR-green master mix (Bioline) in an MJ Opticon thermal cycler, with an initial two-minute denaturation step at 95°C followed by 40 cycles of 95 °C for 5 seconds, 60°C for 10 s, and 72 °C for 10 s. The constitutively expressed gene for glyceraldehyde 3-phosphate dehydrogenase (GAPDH, GiardiaDB GL50803_ 6687) was chosen as an internal reference gene and was amplified with gapdh-F 5’ CCCTTCACGGACTGTGAGTA 3’ and gapdh-R 5’ ATCTCCTCGGGCTTCATAGA 3’ (Barash, 2017). Ct values were determined using the Opticon Monitor software and statistical analyses were conducted using custom Python scripts. All qPCR experiments were performed using two biologically independent samples with four technical replicates. Data are shown as mean expression relative to the GAPDH control sample with 95% confidence intervals.

### Immunostaining of Cas9 and dCas9 CRISPRi knockdown strains

*Giardia* trophozoites were grown to confluency were harvested (see above), washed twice with 6 ml of cold 1X HBS and resuspended in 500 µl of 1X HBS. Cells (250 µl) attached to warm coverslips (37°C, 20 min) were fixed in 4% paraformaldehyde, pH 7.4 (37°C, 15 min), washed three times with 2 ml PEM, pH 6.9 (Woessner and Dawson, 2012), incubated in 0.125M glycine (15 min, 25°C), washed three more times with PEM, and permeabilized with 0.1% Triton X-100 for 10 minutes. After three additional PEM washes, coverslips were blocked in 2 ml PEMBALG (Woessner and Dawson, 2012) for 30 minutes and incubated overnight at 4°C with anti-TAT1 (1:250, Sigma), anti-HA (1:500, Sigma), anti-beta-giardin (1:1000, gift of M. Jenkins, USDA) and/or anti-Cas9 (1:1000, Abcam) antibodies. Coverslips were washed 3 times in PEMBALG and incubated with Alex Fluor 488 goat anti-rabbit and/or Alex Fluor 594 goat anti-mouse antibodies (1:250; Life Technologies) for two hours at room temperature. Coverslips were then washed three times each with PEMBALG and PEM and mounted in Prolong Gold antifade reagent with DAPI (Life Technologies). All imaging experiments were performed with three biologically independent samples.

### Comparing mutant phenotype severity, prevalence, and persistence between CRISPRi and morpholino knockdowns

Morpholino knockdown of median body protein (MBP, GiardiaDB GL50803_16343) was performed as previously described, using the same anti-MBP and mispair morpholino sequences (Woessner and Dawson, 2012). Following electroporation, morpholino knockdown cells were cultured for 24, 48, 72 and 168 hours. The CRISPRi MBP+11 and non-specific gRNA knockdown strains, generated as described above, were cultured from a frozen stock of fully-selected trophozoites passaged for 24, 48, 72 and 168 hours. At each time point, trophozoites were harvested, fixed and stained as described above. Three biologically independent samples were analyzed for each time point and knockdown method.

### Flagellar pair length measurements and ventral disc phenotype analysis

Serial sections of fixed trophozoites were acquired at 0.2 µm intervals using a Leica DMI 6000 wide-field inverted fluorescence microscope with a PlanApo ×100, 1.40 numerical aperture (NA) oil-immersion objective. For flagellar pair length measurements, DIC images were analyzed in FIJI (Schindelin et al., 2012) using a spline-fit line to trace the flagella from the cell body to the flagellar tip. Flagellar length data are presented as mean caudal flagellar length with 95% confidence intervals. For analysis of ventral disc phenotypes, the proportion of aberrant ventral discs and dCas9-positive nuclei within a field of view was determined from maximum intensity projections of processed images. Trophozoites with colocalized DAPI and anti-Cas9 immunostaining were evaluated as positive for dCas9 expression, while cells with undetectable colocalization were scored as dCas9 negative. Colocalization and flagellar length measurements were analyzed, and figures were generated using custom Python scripts.

Super-resolution images of trophozoites exhibiting phenotypes typical of median body protein (MBP, GL50803_16343) silencing by dCas9 were collected at 0.125 µm intervals on a Nikon N-SIM Structured Illumination Super-resolution Microscope with a 100x, 1.49 NA objective, 100 EX V-R diffraction grating, and an Andor iXon3 DU-897E EMCCD. Images were reconstructed in the “Reconstruct Slice” mode and were only used if the reconstruction score was 8. Raw and reconstructed image quality were further assessed using SIMcheck (Ball et al., 2015); only images with adequate scores were used for analysis. Images are displayed as a maximum intensity projection.

## Results

### The SV40 NLS is not sufficient to localize Cas9 to the two Giardia nuclei

To express Cas9 in *Giardia*, we cloned mammalian-codon optimized *Streptococcus pyogenes* Cas9 (mCas9) with a C-terminal SV40 nuclear localization signal (NLS) and a 3XHA epitope tag into the pcGFP1Fpac backbone (Hagen et al., 2011), creating a Cas9-SV40NLS-3XHA-GFP fusion (Figure 1A). A comparison of the modal codon usage frequencies (Davis and Olsen, 2010) of mCas9 and all protein coding genes of *Giardia lamblia* ATCC 50803 indicated that further codon optimization of mCas9 was unnecessary. As Cas9 toxicity has been reported in some systems (Jiang et al., 2014), Cas9 was expressed using a moderate-strength *Giardia* promoter for the constitutive malate dehydrogenase (MDH, GL50803_3331) gene (Figure 1A).

**Figure 1.**
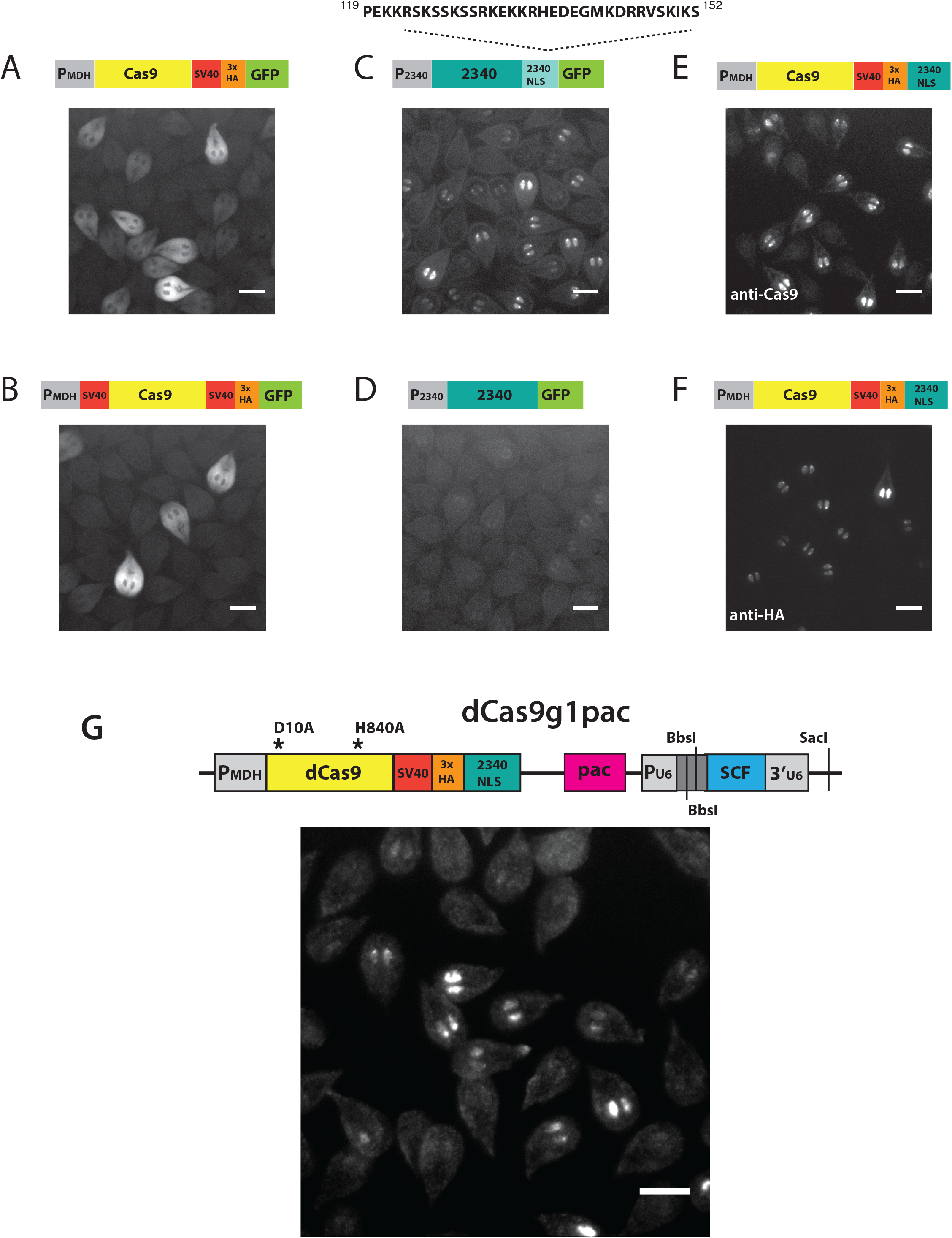
A Giardia-specific nuclear localization signal (NLS) is necessary and sufficient to localize Cas9 and dCas9 to both nuclei. C-terminal GFP-tagged Cas9 with either a single C-terminal SV40 NLS (A) or dual N- and C-terminal SV40 NLSs (B) localizes only to the cytoplasm. The presence of a *Giardia*-specific native 34-amino acid C-terminal NLS from the *Giardia* protein GL50803_2340 (2340NLS) is necessary for the localization of 2340GFP to both nuclei (C, D). The C-terminal addition of the 2340NLS to Cas9 is sufficient for localization to both nuclei as shown by immunostaining with anti-Cas9 (E) or anti-HA (F). A schematic of the *Giardia* CRISPRi vector dCas9g1pac and the localization of dCas9 as determined by anti-Cas9 antibody staining are shown in G. The vector includes catalytically inactive dCas9 with a C-terminal 2340NLS and a 3XHA epitope tag driven by the *Giardia* malate dehydrogenase promoter (P_MDH_), and puromycin resistance marker (pac) for positive selection in *Giardia*. The gRNA cassette is expressed using the *Giardia* U6 spliceosomal RNA pol III promoter, and includes inverted BbsI restriction sites for rapid cloning of specific gRNA target sequences, followed by the gRNA scaffold sequence (SCF). Additional gRNA cassettes are added at the SacI site (Methods). Anti-Cas9 immunostaining indicates over 50% of cells express dCas9 in both nuclei. All scale bars = 5µm.

The Cas9-GFP fusion with a C-terminal SV40 NLS localized only to the cytoplasm, with no signal detected in either of the nuclei (Figure 1A). Cas9 localization to the nucleus is required for Cas9 genome editing, therefore we added a second SV40 NLS at the N-terminus. However, addition of another SV40 NLS had no impact on nuclear localization (Figure 1B). Thus, the SV40 NLS, which is commonly used in other systems and for recombinant Cas9 in commercial Cas9/CRISPR kits, is not sufficient for nuclear localization of Cas9 in *Giardia*. The failure of the SV40 NLS to target full length Cas9 to *Giardia’s* nuclei has been recently noted (Ebneter et al., 2016), although a truncated and inactive Cas9 was successfully localized with this NLS (Ebneter et al., 2016), as were GFP and TetR (Elmendorf et al., 2000).

### Defining a native Giardia NLS to target Cas9 and dCas9 to both nuclei

To localize Cas9 to *Giardia’s* nuclei we used the NLS prediction software NLStradamus to screen for putative NLS sequences (Nguyen Ba et al., 2009). We identified 13 putative NLSs in 65 proteins with known nuclear localization (Supplemental Material). To determine whether the putative NLSs were necessary for nuclear localization, we deleted C-terminal putative NLS sequences from seven nuclear-localizing GFP tagged *Giardia* proteins and assayed for loss of nuclear localization. Deletion of putative NLSs from three proteins resulted in the loss of nuclear localization or an increase in localization to the cytoplasm (Supplemental Material). Deletion of the other four NLSs had no effect on nuclear localization. All three candidate NLSs were sufficient for Cas9 localization to the nuclei (Supplemental Material). Cas9 nuclear localization was the most intense and prevalent with the 34-amino acid C-terminal NLS from the *Giardia* protein GL50803_2340 (Figure C), and deletion of this NLS resulted in loss of localization to the nuclei (Figure 1D). Using the 2340NLS fused to the C-terminus of dCas9 (Figure 1E and F), we created a *Giardia* CRISPR interference expression vector (dCas9g1pac) that included a gRNA cassette with a *Giardia* U6 promoter and puromycin (pac) selection (Methods and Figure 1G).

### CRISPRi-mediated knockdown of the exogenous NanoLuc reporter gene

To assess the ability of CRISPRi to repress transcription in *Giardia*, we created a strain that constitutively expresses the NanoLuc (NLuc) reporter (Hall et al., 2012) from the *Giardia* MDH promoter on a plasmid with neomycin selection. We designed eight gRNAs (+53, +147, +193, +255, +285, +365, +444, +488) (Figure 2A) to target the entire coding region of the NLuc gene at approximately 50 base-pair intervals. One gRNA (-11) was designed to target just upstream of the translation start site. Seven of nine gRNAs significantly repressed NLuc luminescence by ~20-61% as compared to the NanoLucNeo strain (Figure 2B).

**Figure 2.**
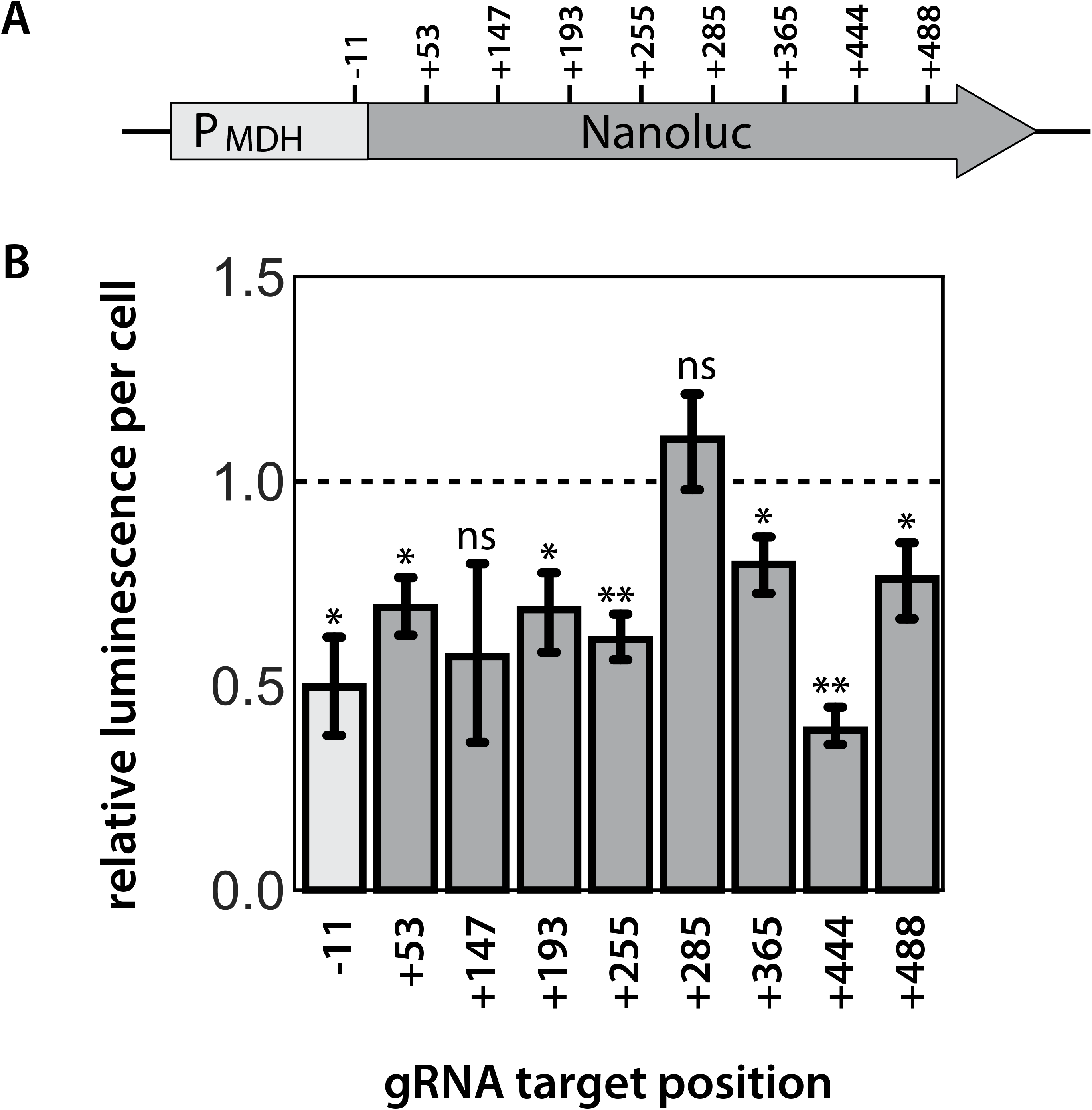
CRISPRi-mediated knockdown of exogenously expressed NanoLuc in Giardia. Nine gRNAs were designed to repress transcription of the luciferase reporter gene NanoLuc expressed exogenously from the *Giardia* MDH promoter (A). Eight gRNAs targeted the coding region of the NanoLuc gene and one gRNA (-11) targeted the region immediately upstream of the translation start site. Target positions are numbered relative to the translational start site ATG and are located 3 bases upstream of the PAM sequence (NGG) for each gRNA. Relative luminescence per cell is plotted for each gRNA designed to repress Nanoluc expression. In B, the mean luminescence per cell for two independent transformations is compared between the different luciferase knockdown strains. Error bars indicate 95% confidence intervals. The significance of luciferase knockdown relative to a non-specific gRNA control was assessed using unpaired t-test, with ns = not significant, *p ≤ 0.05, **≤0.01 (Methods).

### CRISPRi mediated knockdowns of kinesin-13 or kinesin-2a cause significant alterations in flagellar length

*Giardia* has four pairs of bilaterally symmetric flagella that are maintained at consistent equilibrium lengths (Figure 3A). *Giardia* possesses a single heterotrimeric kinesin-2 motor, consisting of kinesin-2a, 2b, and the kinesin-associated protein (KAP) that is required for flagellar assembly and maintenance (Hoeng et al., 2008). The length of a flagellum is determined by the balance of kinesin-2 dependent assembly (Kozminski et al., 1995) and kinesin-13 dependent disassembly at the distal flagellar tip (Dawson et al., 2007). The dominant-negative overexpression of a rigor mutant kinesin-13 causes significant increases in the length of *Giardia’s* eight flagella, larger median bodies, and cell division defects (Dawson et al., 2007). In contrast, both morpholino-based knockdown of kinesin-2b and dominant negative expression of a rigor mutant kinesin-2a result in significant flagellar shortening and mitotic defects (Hoeng et al., 2008).

**Figure 3.**
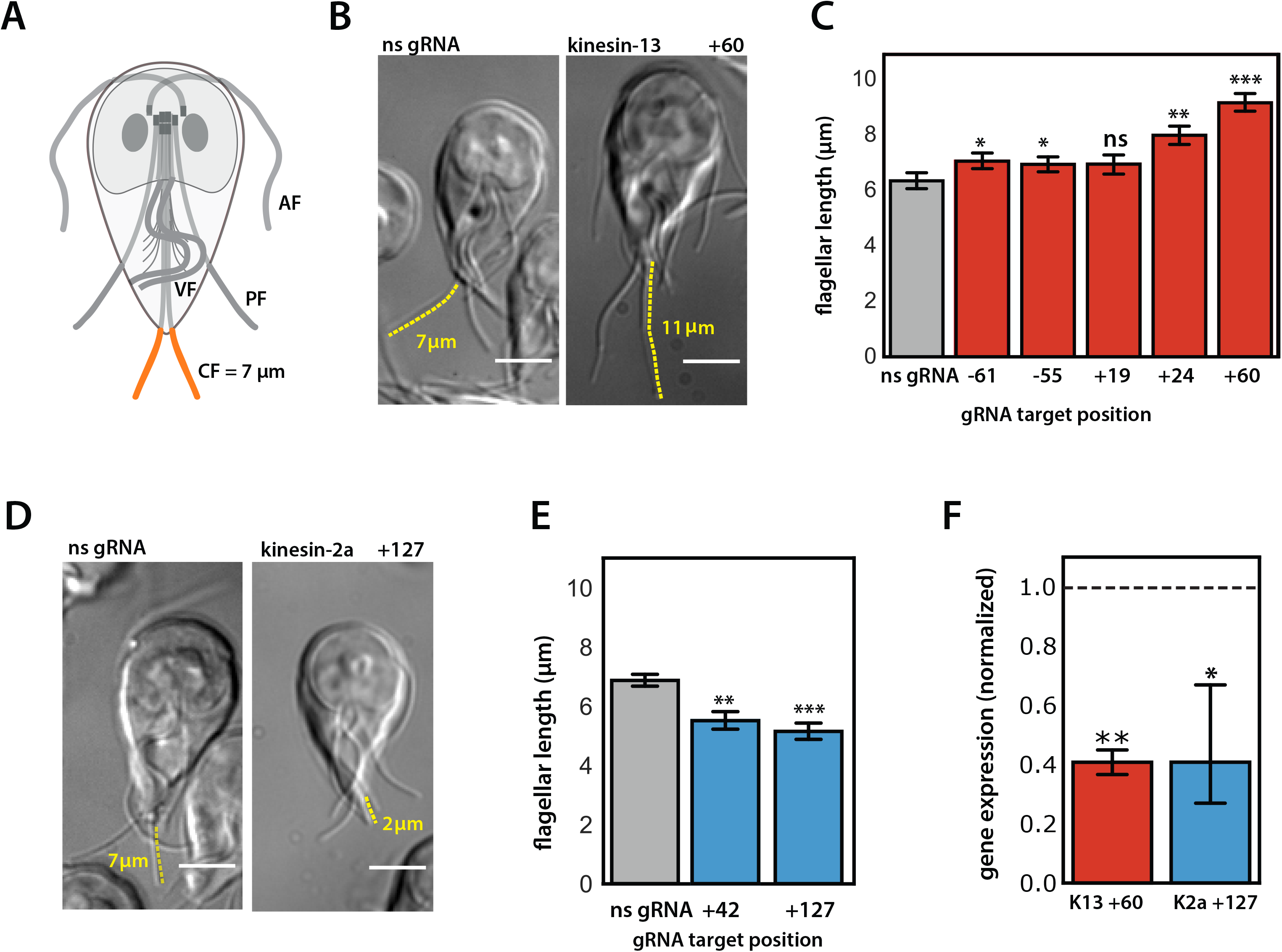
CRISPRi mediated knockdowns of kinesin-13 or kinesin-2a cause significant alterations in flagellar length. *Giardia* has four pairs of bilaterally symmetric flagella – anterior (AF), ventral (VF), posteriolateral (PF), and caudal (CF, orange) – with distinct equilibrium interphase lengths (A). Representative images and quantification of caudal flagellar length are shown for the CRISPRi-mediated knockdown of two endogenous kinesins known to regulate flagellar length in *Giardia*: kinesin-13 +60 gRNA (B) and kinesin-2a +127 gRNA (D) as compared to a non-specific gRNA strain (B and D, see Methods). Yellow traces highlight representative caudal flagella for flagellar length measurements. Scale bars = 5 µm. In C, average caudal flagellar length for five kinesin-13 knockdown strains with gRNAs targeting the upstream region (-61, -55) or coding region (+19, +24, and +60) are compared to the strain expressing a non-specific gRNA (ns gRNA). In E, the average caudal flagellar length is quantified for strains with gRNAs targeting +42 and +127 positions of kinesin-2a as compared to a non-specific gRNA expressing strain (ns gRNA). All images were acquired from three independent experiments, with more than 70 cells analyzed for each treatment group. In F, the decrease in gene expression of kinesin 13 (red) in the kinesin-13 +60 strain and kinesin-2a (blue) in the kinesin-2a +127 strain is shown relative to expression levels of the two kinesins in a non-specific gRNA expressing strain (dotted line indicates expression in the ns gRNA strain). Kinesin gene expression is normalized to the *Giardia* GAPDH gene from two independent experiments. In all panels, error bars indicate 95% confidence intervals, and significance was assessed using unpaired t-test with ns = not significant, *p≤0.05, **≤0.01, ***≤0.001.

To determine the ability of CRISPRi to knockdown endogenous genes in *Giardia*, we designed five gRNAs (-61, -55, +19, +24, +60) to target kinesin-13 (GiardiaDB GL50803_16945) and two gRNAs (+42, +127) to target kinesin-2a (GiardiaDB GL50803_17333). To assess the flagellar length defects of kinesin-13 and kinesin-2a knockdowns, we fixed cells and measured the length of the caudal flagella, a convenient reporter for flagellar length (Dawson et al., 2007). Four of five gRNAs targeting kinesin-13 (-61, -55, +24, +60) significantly increased the length of caudal flagella compared to a non-specific gRNA (Figure 3B,C). While both gRNAs targeting the promoter region of kinesin-13 (-61, -55) resulted in significant length increases for caudal flagella, the length defects were less severe than for the two gRNAs (+24, +60) targeting the coding region (Figure 3C). Both gRNAs targeting kinesin-2a (+42, +127) significantly decreased caudal flagellar length compared to a non-specific gRNA (Figure 3D,E).

To determine the degree of transcriptional repression of kinesin-13 and kinesin-2a conferred by CRISPRi knockdown, we quantified the expression of these targets in the kinesin-13 +60 and kinesin-2a +127 strains using quantitative PCR (qPCR). Expression of either target was reduced by 59% as compared to a non-specific gRNA strain (Figure 3F).

### CRISPRi knockdown of MBP causes severe disc defects that are highly prevalent and persist at least one week in cultured trophozoites

Attachment to the host intestinal epithelium by the ventral disc – a highly ordered and complex microtubule (MT) array – is critical to *Giardia*’s pathogenesis in the host (Nosala et al., 2018). The ventral disc is defined by more than 90 parallel and uniformly spaced MTs that spiral into a circular domed structure (Crossley and Holberton, 1983; Crossley and Holberton, 1985; Feely et al., 1982; Friend, 1966; Holberton, 1973; Holberton, 1981). MBP (GiardiaDB GL50803_16343) is one of 87 proteins that localize to the ventral disc (Nosala et al., 2018), and morpholino knockdown of MBP results in ventral discs with an “open” and “flattened” conformation, as well as decreased attachment efficiency (Woessner and Dawson, 2012).

To test the efficacy of MBP knockdown using CRISPRi, we created three strains with gRNAs targeting either the promoter (-7) or coding regions (+11, +1566) of the gene. The proportions of aberrant ventral disc phenotypes in each knockdown strain were determined by fixing and immunostaining trophozoites for dCas9 and the ventral disc protein β-giardin (Baker et al., 1988). Images were scored for disc phenotype and dCas9 expression in each cell (Figure 4A). In each MBP knockdown strain, the presence of an aberrant ventral disc was positively correlated with dCas9 nuclear staining (Figure 4A,C). Trophozoites positive for dCas9 nuclear staining had severe ventral disc structural defects with incompletely closed (or aberrant) discs (Figure 4A,B) as compared to the typical closed, domed disc structure (Woessner and Dawson, 2012). Specifically, 46-93% of trophozoites with dCas9 nuclear staining had aberrant ventral disc structures using any of the three gRNAs (Figure 4C). Ventral discs in the non-specific gRNA strain lacked structural defects and were comparable to wild-type. Super resolution imaging using structured illumination microscopy (SIM) highlights the disrupted and open ventral disc ultrastructure in the CRISPRi knockdowns (Figure 4B), with some dCas9 positive cells lacking large regions of the ventral disc (Figure 4B).

**Figure 4.**
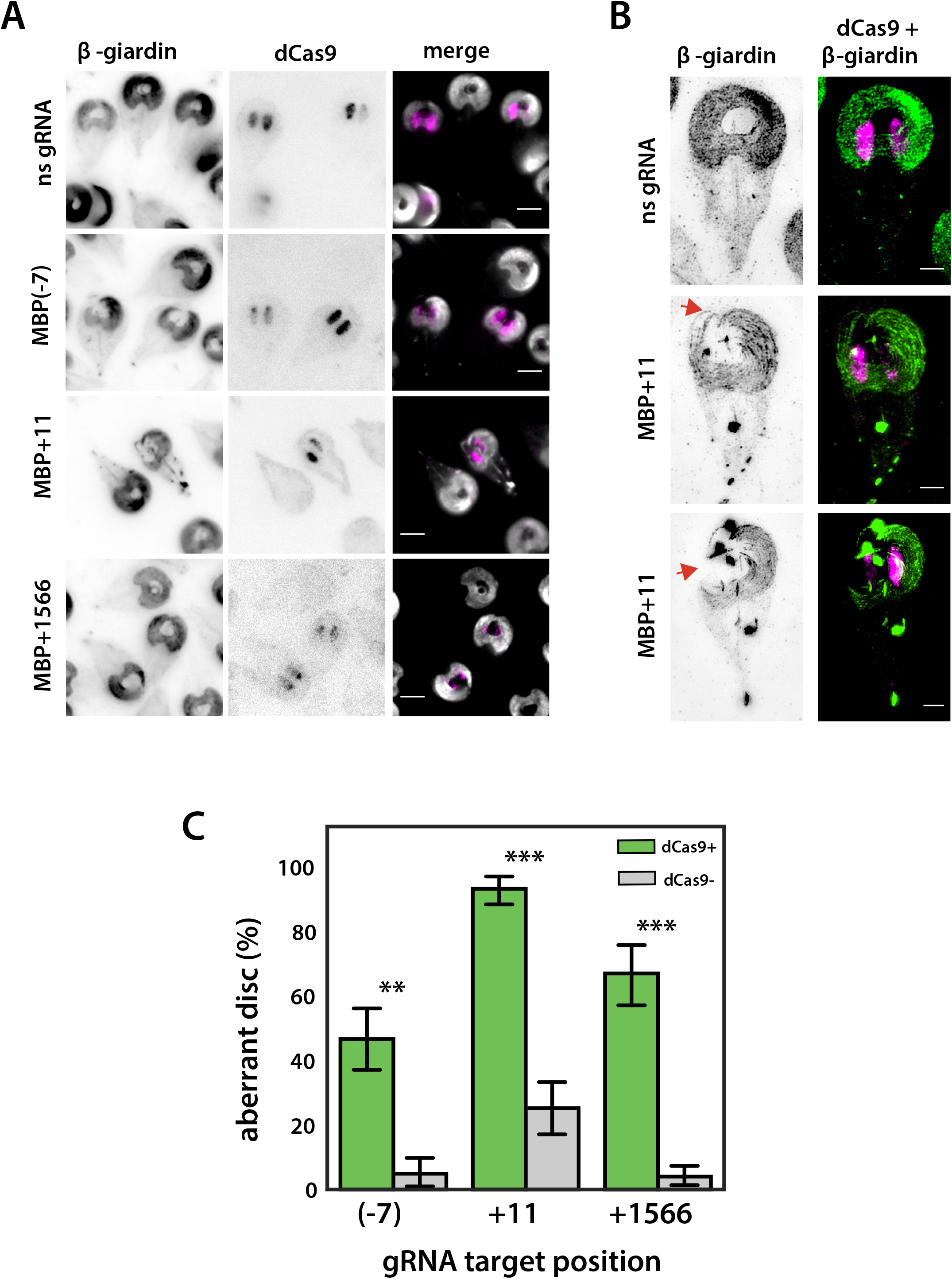
CRISPRi knockdown of the ventral disc protein MBP causes severe structural defects. In A, widefield images are presented showing immunostaining of both the disc protein β-giardin (left) and dCas9 (middle) in strains expressing a non-specific gRNA, or CRISPRi gRNAs designed to knock down the median body protein (MBP(-7) gRNA, MBP+11 gRNA, and MBP+1566 gRNA). Scale bars are 5 µm. In B, representative structured illumination microscopy (SIM) images of the MBP+11 gRNA strain immunostained for β-giardin (left) and dCas9 (right) highlight the characteristic “open” disc conformation (arrows) observed with morpholino knockdown of MBP ((Woessner and Dawson, 2012), and see Figure 5B below). Scale bar = 2 µm. In C, the proportion of aberrant disc phenotypes in strains expressing MBP(-7), MBP+11, and MBP+1566 gRNAs is quantified and compared in cells scored as dCas9 positive (green) or negative (gray). Three independent experiments were performed, with more than 200 cells analyzed for each gRNA. Error bars indicate 95% confidence intervals. Significance was assessed using unpaired t-test with **p ≤0.01 and ***≤0.001.

**Figure 5.**
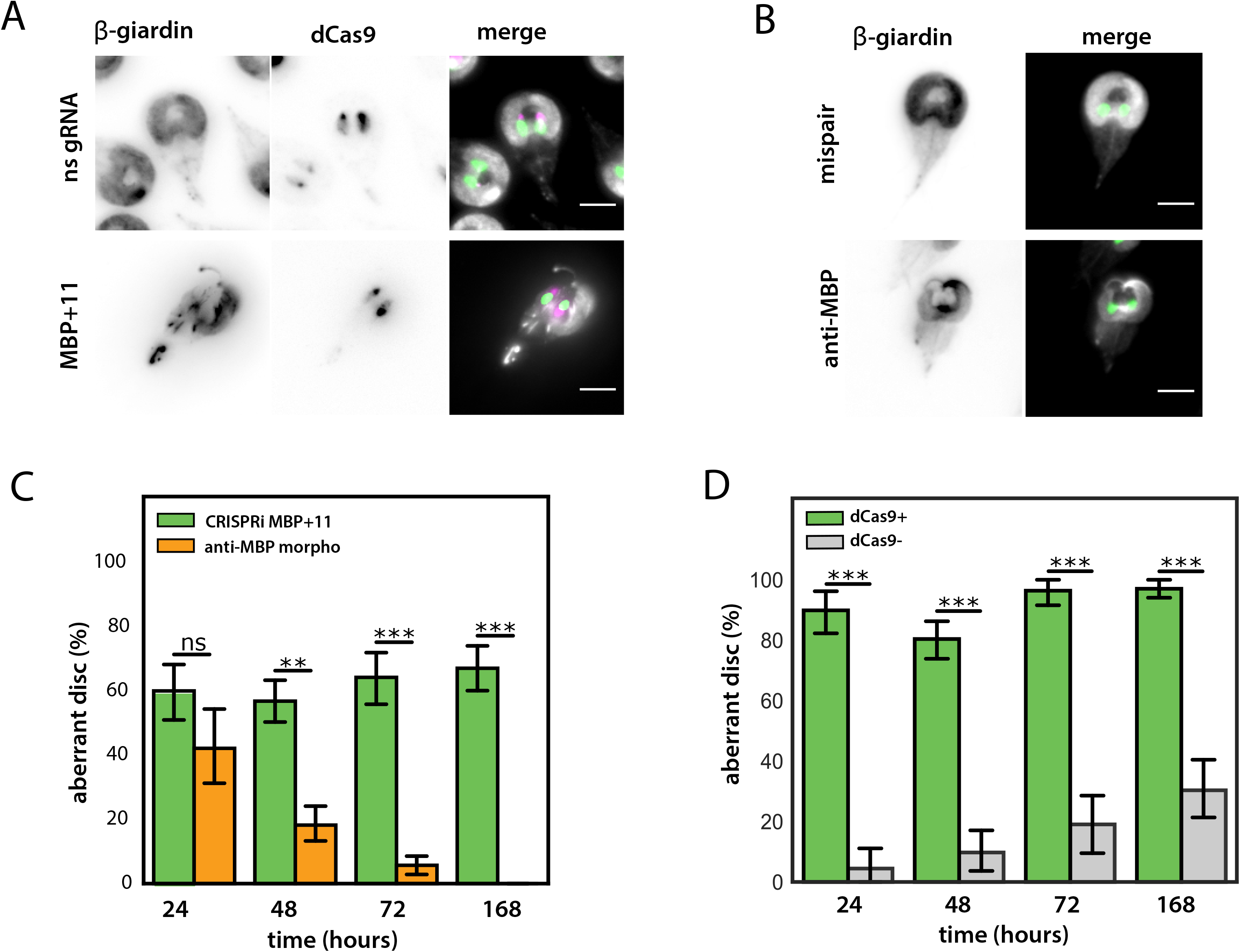
Aberrant disc phenotypes in the CRISPRi MBP+11 knockdown strain are highly penetrant and stable. The open disc phenotype of the MBP+11 gRNA strain (A) is similar to aberrant disc phenotypes observed using an anti-MBP morpholino (B, and see (Woessner and Dawson, 2012)). DAPI = green, anti-Cas9 = pink. Scale bar = 5 µm. Unlike the anti-MBP morpholino knockdown (orange), the open disc phenotype of the CRISPRi MBP+11 strain (green) is stable in cultured cells for at least one week (C). The proportion of dCas9+ (green) cells with the open disc phenotype is also consistent and highly penetrant in the MBP+11 strain (D). Three independent experiments were performed, with more than 75 cells analyzed for each gRNA. Error bars indicate 95% confidence intervals. Significance was assessed using unpaired t-test with ns = not significant, **p ≤0.01 and ***p≤0.001.

To assess the stability and persistence of CRISPRi knockdown phenotypes, we compared the proportion of trophozoites with aberrant ventral discs obtained via the CRISPRi knockdown to those obtained via morpholino knockdown of MBP over the course of one week. Translational knockdown with morpholinos is currently the only knockdown method widely used in *Giardia*, yet morpholinos are costly and only transiently repress protein expression (Carpenter and Cande, 2009).

To directly compare the severity, penetrance, and persistence of aberrant discs after MBP knockdown using CRISPRi or morpholinos, we fixed and immunostained trophozoites cultured for 24, 48, 72, and 168 hours after introduction of an anti-MBP morpholino and compared them to the constitutively expressed CRISPRi MBP+11 strain. The proportion of cells with aberrant discs was comparable for both methods at 24 hours. With prolonged passage in culture, however, the penetrance of aberrant discs phenotypes in the morpholino knockdown population rapidly decreased, with significantly fewer aberrant discs present at 48, 72, and 168 hours (Figure 5C). At 168 hours, no aberrant disc phenotypes were observed in the morpholino knockdown population. In contrast, the penetrance (e.g., the proportion of aberrant disc phenotypes in the CRISPRi strain) remained close to 60% of total cells for 168 hours (Figure 5C). Furthermore, more than 85% of dCas9 positive cells had aberrant disc phenotypes at 24 hours, and this high degree of penetrance persisted to 168 hours of passage in culture (Figure 5D).

### Simultaneous knockdown of kinesin-13 and MBP results in trophozoites with prevalent mutant phenotypes

Simultaneous repression of multiple genes is commonly used to interrogate epistatic interactions or functional redundancy of paralogous genes. The flexibility of the CRISPR/Cas9 system we designed for *Giardia* permits the simultaneous targeting of multiple regions of the same target or of multiple target genes by simply including multiple gRNAs. To test these versatile aspects of CRISPRi in *Giardia*, we modified the dCas9g1pac vector to simultaneously express two different gRNAs. In short, we designed a set of primers to amplify the entire gRNA expression cassette from an existing gRNA vector and added this amplified region downstream of the gRNA cassette of a second gRNA vector (Figure 6A). Specifically, we added the MBP+11 gRNA cassette to the kinesin-13+60 vector, as each of these gRNAs provided strong repression of their respective targets with distinct phenotypes (Figures 3C, 4B).

**Figure 6.**
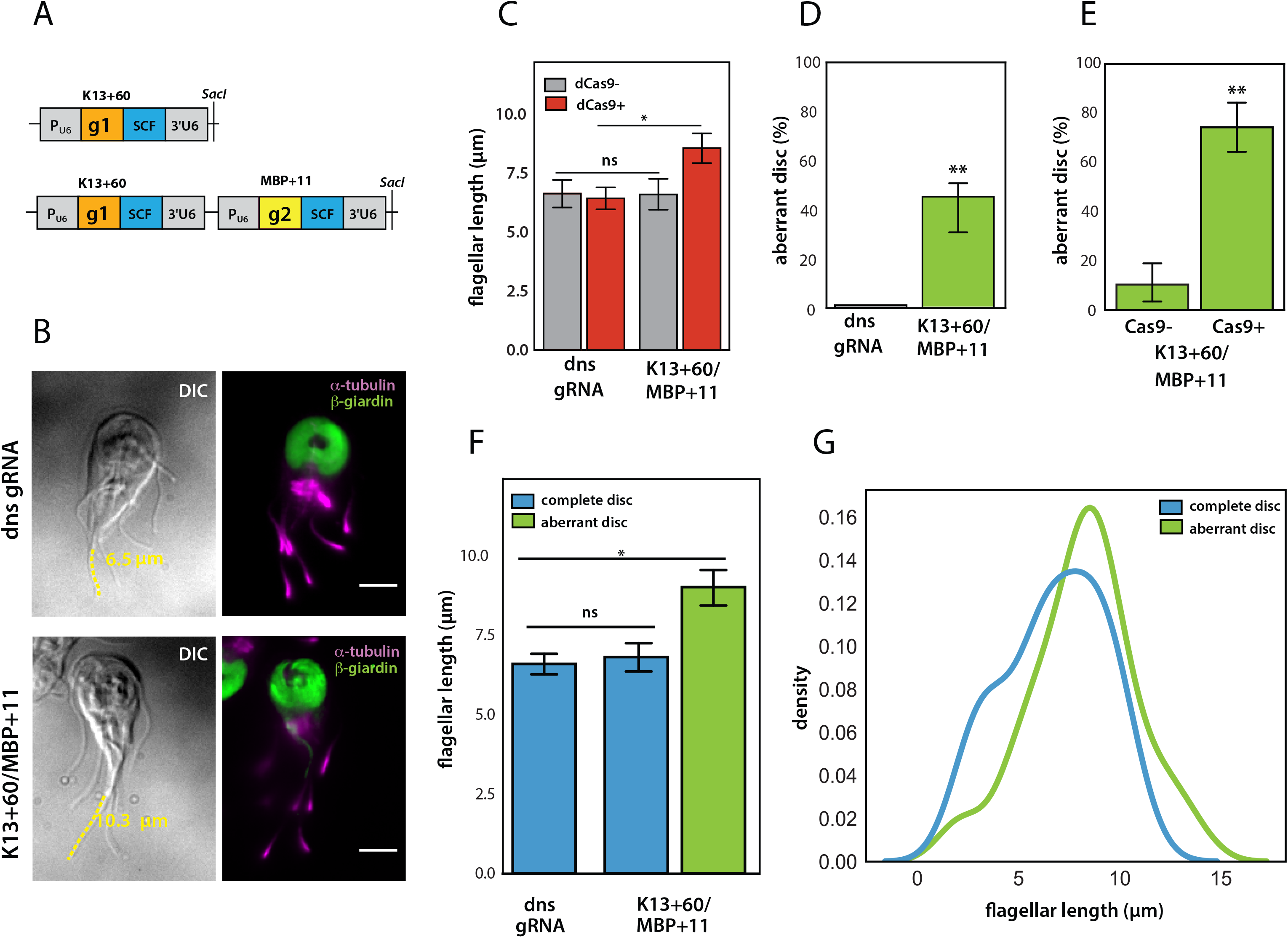
Simultaneous knockdown of kinesin-13 and MBP using two gRNAs to target both genes. For simultaneous expression of two gRNAs targeting both MBP and kinesin-13, we cloned the two gRNA cassettes g1 (MBP+11) and g2 (kinesin-13+60 in tandem. Scaffold sequence = SCF (A). The gRNA cassette is expressed using the *Giardia* U6 spliceosomal RNA pol III promoter (P_U6_). In B, representative images for the knockdown of both kinesin-13 and MBP in a strain expressing both kinesin-13+60 and MBP+11 gRNAs highlight a cell with both the open disc and long flagella phenotypes as compared to a cell expressing the dual non-specific gRNAs (dns gRNA). dCas9 positive (red) cells of the K13+60/MBP+11 dual knockdown strain had significantly longer caudal flagella as compared to dCas9 negative cells or those expressing the non-specific gRNAs (C). Aberrant disc phenotypes (green) were observed in over 45% of cells in the K13+60/MBP+11 dual knockdown strain (D), with high penetrance of aberrant disc phenotypes in dCas9+ cells (E). Cells with aberrant discs (green) also had significantly longer caudal flagella than dns gRNA or K13+60/MBP+11 cells with complete discs (blue) (F). Shifts in the caudal flagellar length distributions (represented using kernel density estimates) show that cells with long flagella have open disc phenotypes (G). Two independent experiments were performed, with more than 95 cells analyzed for each gRNA. Error bars indicate 95% confidence intervals. Significance was assessed using unpaired t-test with ns = not significant, *p ≤0.01 and **p≤0.001.

To assess the efficacy of targeting multiple genes with CRISPRi, we fixed and stained trophozoites expressing dCas9 and the kinesin-13+60 and MBP+11 gRNAs. Flagellar and disc phenotypes were compared to a dCas9 ‘dual’ non-specific gRNA (dns gRNA) control strain that expressed the non-specific gRNA sequence from two gRNA cassettes. We quantified flagellar length and the presence of open discs within the same cell and found that a high proportion of the kinesin-13+60/MBP+11 cells had both mutant phenotypes (Figure 6B).

Caudal flagella were 16% longer in the kinesin-13+60/MBP+11 knockdown strain as compared to the dual non-specific gRNA strain. Like the single MBP+11 strain (Figure 3C), trophozoites in the kinesin-13+60/MBP+11 knockdown strain that were positive for dCas9 staining had more severe defects, with caudal flagella that were 30% longer than in cells lacking dCas9 staining (Figure 6C). With respect to ventral disc phenotypes, 45% of kinesin-13+60/MBP+11 knockdown trophozoites had an aberrant “open” ventral disc structure, while no ventral discs in the dual non-specific gRNA strain had structural defects (Figure 6D). As with caudal flagellar length defects, open or aberrant ventral disc phenotypes were highly prevalent in dCas9 positive cells; 74% of dCas9 positive cells had aberrant ventral discs as compared to only 10% of dCas9 negative cells (Figure 6E). Lastly, trophozoites with aberrant discs were primarily found in the fraction of cells with longer caudal flagella (Figure 6F), and especially in cells with caudal flagellar lengths greater than 10 µm (Figure 6G).

## Discussion

CRISPRi is a robust alternative to RNAi-mediated gene silencing for precise knockdown of gene expression in both eukaryotic and bacterial model systems (Kampmann, 2018; Larson et al., 2013). Our successful use of CRISPRi to repress both exogenous (Figure 2) and single or multiple endogenous genes (Figures 3-6) in *Giardia* underscores the versatility of this stable, modular, and efficient gene regulation system. CRISPRi-based gene repression will rapidly change how we study basic cell biology, development, and pathogenesis in this widespread and understudied binucleate parasite.

### Development of CRISPRi-mediated knockdown in Giardia

Molecular genetic tool development in *Giardia* has focused on transient translational repression by electroporation of morpholinos (Carpenter and Cande, 2009) or on the overexpression of long double-stranded RNAs or hammerhead ribozymes for transcriptional repression (Chen et al., 2007; Dan et al., 2000). While CRISPR/Cas9-mediated knockout strategies have recently been used for genome engineering in several parasitic protists (Ren and Gupta, 2017), this is the first demonstration of CRISPRi-mediated transcriptional repression in these organisms.

The *Giardia* CRISPRi expression system is compact and self-contained on a single episomal plasmid (Figure 1G). Expression of the modular dCas9 and gRNA CRISPRi cassettes does not require *Giardia* host or viral factors as is required for antisense (Rivero et al., 2010) or hammerhead ribozyme-mediated transcriptional repression (Chen et al., 2007; Dan et al., 2000). A native *Giardia* NLS is required for targeting of the Cas9 or dCas9/gRNA DNA recognition complex to both nuclei (Figure 1E-G). Once imported into the nuclei, the Cas9/gRNA complex is targeted to a specific genomic locus where it sterically interferes with RNA polymerase or transcription factor binding, or with transcriptional elongation, as has been shown in bacteria or other eukaryotes (Larson et al., 2013). The ability to direct dCas9 or other native transcriptional elements to both nuclei in *Giardia* will enable the modulation of transcriptional networks not only by repression, but also by differential expression through the fusion of dCas9 to *Giardia*-specific transcription factors (Kampmann, 2018).

The choice of gRNA target site impacts the degree of transcriptional repression in every CRISPRi system (Larson et al., 2013). In both bacteria and human cell lines, targeting gRNAs close to the translation start site results in stronger repression (Gilbert et al., 2013; Qi et al., 2013). We systematically targeted eight sites within the coding region of exogenously expressed NanoLuc (NLuc), a luminescent reporter gene, to determine how gRNA positioning might influence the magnitude of transcriptional repression in *Giardia* (Figure 2).

Guide RNAs targeting all regions of the NanoLuc reporter substantially repressed luminescence; however, there was no correlation between gRNA target site position and the magnitude of repression. Because we used a native *Giardia* promoter (MDH) for NanoLuc expression, we could not target this region without potentially altering expression of the native MDH gene. This limits our interpretation of knockdown results for the gRNA that targeted the region immediately upstream of NanoLuc. For the kinesin-13 and MBP CRISPRi knockdowns, however, significant transcriptional repression and aberrant phenotypes were observed for gRNAs targeting regions both upstream and downstream of the translation start site (Figure 3 and Figure 4). More robust and persistent phenotypes for kinesin-2a, kinesin-13, and MBP were associated with gRNAs that targeted the coding regions of these genes, which is consistent with the inhibition of transcriptional elongation, rather than inhibition of transcriptional initiation (Larson et al., 2013). Furthermore, due to the ill-defined nature of *Giardia* promoters and the limited length of intergenic regions (Davis-Hayman and Nash, 2002), we recommend that several gRNAs be designed to target the coding region for successful repression in *Giardia*.

### Highly penetrant and persistent transcriptional repression of endogenous genes

Using the eight flagella, motile *Giardia* trophozoites colonize the upper gastrointestinal tract (Dawson and House, 2010b), attaching extracellularly to the intestinal villi using the ventral disc, thereby resisting peristaltic flow (Elmendorf et al., 2003; Nosala and Dawson, 2015). Our demonstration of efficient and highly penetrant CRISPRi-mediated transcriptional repression of three endogenous cytoskeletal proteins (Figures 3-6) highlights the versatility of this method to interrogate the contribution of the cytoskeleton to parasite motility and attachment.

Each of the CRISPRi-mediated knockdowns of endogenous genes resulted in cytoskeletal phenotypes that were consistent with phenotypes observed in *Giardia* in prior studies (Carpenter and Cande, 2009; Dawson et al., 2007; Hoeng et al., 2008; Woessner and Dawson, 2012). In *Giardia* and other flagellates, alterations in flagellar assembly or disassembly dynamics cause flagellar length defects (Carpenter and Cande, 2009; Dawson et al., 2007; Hoeng et al., 2008; Sloboda, 2005; Woessner and Dawson, 2012). Both electroporation of anti-kinesin-2b morpholinos (Carpenter and Cande, 2009), and inducible dominant negative overexpression of kinesin-2a (Hoeng et al., 2008) limit the degree of IFT-mediated assembly, resulting in shorter flagella in *Giardia*. In contrast, overexpression of a dominant negative kinesin-13 results in increased length of all eight flagella by decreasing the rate of disassembly (Dawson et al., 2007; Hoeng et al., 2008). When we knocked down these kinesins using CRISPRi with gRNAs targeting upstream or downstream of the translation start site, we observed flagellar length increases (up to 60% longer for kinesin-13, Figure 3B,C) or decreases (up to 30% shorter for kinesin-2a, Figure 3D,E). These length variations are comparable to those seen with morpholino knockdowns or overexpression of dominant negatives for kinesin-13 or kinesin-2a (Dawson et al., 2007; Hoeng et al., 2008), underscoring the evolutionarily conserved and essential roles these two kinesins in flagellar length regulation in *Giardia*.

*Giardia*’s ventral disc is essential for attachment to the host, and is composed of nearly 90 proteins (Nosala et al., 2018). The CRISPRi-mediated knockdown of the disc-associated protein median body protein (MBP) also confirms the same “open” and “flat” disc phenotype (Figure 4) observed using transient morpholino translational knockdowns (Woessner and Dawson, 2012) (Figure 5A,B). Similar to prior morpholino knockdowns, we also saw extreme defects in ventral disc structure, including partial discs that were most evident with the gRNA targeting the +11 position of the MBP coding region (Figure 4B).

The overall penetrance and stability of CRISPRi knockdown mutant phenotypes in a population is critical for evaluating the efficacy of the *Giardia* CRISPRi knockdown system. For CRISPRi MBP knockdown with any of three gRNAs, the degree of dCas9 expression was highly correlated with aberrant disc structure (Figure 4C). The penetrance of aberrant disc phenotypes was 46% to 93% in the population of dCas9 positive cells as compared to 4% to 25% penetrance in dCas9 negative cells. Thus, in addition to testing multiple candidate gRNAs for severity of phenotypes, we also advocate the use of FACS or similar methods of cell sorting to enrich for a more homogeneous population expressing dCas9, as has been done in other CRISPRi systems (Gilbert et al., 2013).

The stability or persistence of the CRISPRi knockdown phenotype in a population over many generations is also a key feature of the *Giardia* CRISPRi system, which includes a positive selectable marker (pac) to allow maintenance of the CRISPRi plasmid under puromycin selection. By passaging the MPB+11 strain repeatedly for one week, we showed that the penetrance of the aberrant disc phenotype remained close to 60% of total cells for up to 168 hours (Figure 5C). Between 80% to 97% of dCas9 positive cells had open, incomplete ventral discs at any time point up to 168 hours in culture (Figure 5D). Thus, CRISPRi knockdowns are not only highly penetrant but also highly stable as compared to transient knockdowns with morpholinos, wherein the prevalence of cells with aberrant discs rapidly decreased in the population after 48 hours and was completely lost by 168 hours (Figure 5).

As compared to transient and costly morpholino knockdowns, CRISPRi produces stable knockdown strains that are positively selected and can be archived and evaluated at any time. The degree of morpholino knockdown depends on the initial transformation efficiency, and morpholino knockdowns require analysis immediately after electroporation, which could introduce phenotypic artefacts or variability. In contrast, multiple CRISPRi knockdown strains can be made using one or more gRNAs that target the same gene, permitting a comparison of mutant phenotypes in different knockdown strains (Figures 2-5). Furthermore, the generation of CRISPRi knockdown vectors can be multiplexed and only requires the purchase and annealing of two short oligomers.

### Knockdown of multiple genes by the simultaneous expression of multiple gRNAs

The modular design of the *Giardia* gRNA expression cassette allows for concatenation of two or more gRNAs to target multiple sites on a single gene, or the targeting of more than one *Giardia* gene for phenotypic analysis (Figure 6). Simultaneous CRISPRi knockdown of both the flagellar length regulator kinesin-13 and the disc-associated protein MBP resulted in highly penetrant flagellar length and disc defects (Figure 6). Furthermore, the majority of cells with flagellar length defects also had aberrant discs (Figure 6F), and the cells with the longest flagellar lengths were exclusively those with open, incomplete discs. Thus, the ability to create stable CRISPRi knockdown strains with defects in multiple genes now allows the evaluation of the functional redundancy of similar and paralogous genes in *Giardia*, as well as the interrogation of epistatic interactions of genes in a biochemical pathway.

### Using CRISPRi to identify and evaluate genes critical in giardiasis

Over 40 percent of *Giardia’s* 6000 genes encode “hypothetical” proteins that lack similarity to proteins in the human host (Morrison et al., 2007). Many hypothetical proteins are highly expressed during *in vivo* infections yet lack any known cellular function (Pham et al., 2017). Rapid and stable CRISPRi knockdown enables the functional evaluation of such hypothetical genes as druggable targets for giardiasis. Combining stable CRISPRi knockdowns with our recently developed bioluminescent imaging (BLI) to monitor temporal and spatial patterns of *Giardia* infection dynamics and metabolism in the host (Barash, 2017) will allow the examination of *Giardia* mutants with respect to *in vivo* fitness, colonization, or encystation defects.

The lack of forward genetic tools for *Giardia* has limited our ability to define genes that are required for basic parasite biology. CRISPRi knockdowns with partial transcriptional repression facilitate the identification of genes with severe fitness costs, as the complete knockdown or knockout of essential genes results in lethal phenotypes. As compared to reverse genetic approaches, the use of untargeted, genome-wide CRISPRi screens (Kampmann, 2018; Larson et al., 2013) could identify essential genes critical for *Giardia* growth, differentiation, and pathogenesis.

## Accession Numbers

The *Giardia* CRISPRi and NanoLuc reporter vectors have been deposited in GenBank with the following accession numbers: dCas9g1pac (MH037009), NanoLucNeo (MH037010).

## Acknowledgements

This work was supported by NIH/NIAID awards R01AI077571 and R21AI119791-01 to SCD. Plasmids JDS246 (Addgene #43861) and MSP712 (Addgene #65768) were gifts from Keith Joung. Plasmid pKS_mNeonGreen-N11_NEO was a gift from Alex Paredez (University of Washington, Seattle). The anti-beta-giardin antibody was a gift of Mark Jenkins (USDA, ARS, Animal Parasitic Diseases Laboratory). We thank the MCB Light Microscopy Imaging Facility, a UC Davis Campus Core Research Facility, for the use of the Nikon N-SIM Structured Illumination Super-resolution Microscope.

